# 116 independent genetic variants influence the neuroticism personality trait in over 329,000 UK Biobank individuals

**DOI:** 10.1101/168906

**Authors:** Michelle Luciano, Saskia P Hagenaars, Gail Davies, W David Hill, Toni-Kim Clarke, Masoud Shirali, Riccardo E Marioni, Sarah E Harris, David C Liewald, Chloe Fawns-Ritchie, Mark J Adams, David M Howard, Cathryn M Lewis, Catharine R. Gale, Andrew M McIntosh, Ian J Deary

**Affiliations:** Centre for Cognitive Ageing and Cognitive Epidemiology, Department of Psychology, School of Philosophy, Psychology and Language Sciences, The University of Edinburgh, Edinburgh, UK; MRC Social, Genetic and Developmental Psychiatry Centre, Institute of Psychiatry, Psychology & Neuroscience, King’s College London, De Crespigny Park, Denmark Hill, London, SE5 8AF, UK; Division of Psychiatry, University of Edinburgh, Edinburgh, UK; Centre for Genomic & Experimental Medicine, MRC Institute of Genetics & Molecular Medicine, University of Edinburgh, Western General Hospital, Edinburgh, UK; MRC Lifecourse Epidemiology Unit, University of Southampton, Southampton, UK.

**Author notes:** Corresponding author, Department of Psychology, The University of Edinburgh, 7 George Square, EH8 9JZ, Edinburgh, UK.

## Abstract

Neuroticism is a stable personality trait ^1^; twin studies report heritability between 30% and 50% ^2^, and SNP-based heritability is about 15% ^3^. Higher levels of neuroticism are associated with poorer mental and physical health ^4,5^, and the economic burden of neuroticism for societies is high ^6^. To date, genome-wide association (GWA) studies of neuroticism have identified up to 11 genetic loci ^3,7^. Here we report 116 significant independent genetic loci from a GWA of neuroticism in 329,821 UK Biobank participants, with replication available in a GWA meta-analysis of neuroticism in 122,867 individuals. Genetic signals for neuroticism were enriched in neuronal genesis and differentiation pathways, and substantial genetic correlations were found between neuroticism and depressive symptoms (r_g_ = .82, SE=.03), major depressive disorder (r_g_ = .69, SE=.07) and subjective wellbeing (rg = -.68, SE=.03) alongside other mental health traits. These discoveries significantly advance our understanding of neuroticism and its association with major depressive disorder.

---

Understanding why people differ in neuroticism will be an important contribution to understanding people’s liability to poor mental health through the life course. The strong genetic correlation between neuroticism and mental health, especially anxiety and major depressive disorder ^8,9^, means that exploring the genetic contribution to differences in neuroticism is one way to understand more about these common and burdensome, but aetiologically intractable illnesses. In the largest GWA study of major depression, 44 independent genetic loci have been identified ^10^.

The UK Biobank open resource (**http://www.ukbiobank.ac.uk**) has health and medical information for over 500,000 individuals aged 40-69 years from the United Kingdom, assessed between 2006 and 2010 ^11^; genetic data are also available ^12^. We performed a GWA analysis of trait neuroticism in 329,821 unrelated adults (152,710/46.3% male) of European descent from this resource who had high-quality genotype data (Online Methods). Neuroticism was measured by the total score of the 12-item Eysenck Personality Questionnaire-Revised Short Form (EPQ-R-S) ^13^ (Supplementary Table 1 and Supplementary Fig. 1; Online Methods). The score used in the association analysis was residualized for the effects of age, sex, assessment centre, genotype batch, array, and 40 genetic principal components. Neuroticism scores were tested against 18,485,882 bi-allelic single nucleotide polymorphism (SNP) variants, based on the Haplotype Reference Consortium panel, with a minor allele frequency ≥ 0.0005 and an information/imputation quality score of ≥ 0.1 under an additive model using the BGENIE software ^12^. The distribution of obtained versus expected results under the null hypothesis showed some genomic inflation, with a lambda of 1.15 (the quantile-quantile plot is shown in Supplementary Fig. 2). Using univariate linkage disequilibrium score regression (LDSR) ^14^, 3.2% of this inflation was shown to be due to the presence of a large polygenic signal, indicated by the intercept being close to 1 (1.02, SE = .01). SNP-based heritability of neuroticism was calculated using LSDR, and was estimated at .108 (SE=.005).

Genome-wide significance (P < 5 × 10^−8^) was demonstrated for 10,353 genetic variants with a further 17,668 variants supported at a suggestive level (P < 1 × 10^−5^) of significance (Supplementary Table 3). The Manhattan plot is shown in Figure 1 and gene annotation for the significant 1000G SNPs is shown in Supplementary Table 4. Using the PLINK clumping tool ^15^, 116 of the significant SNPs were shown to be independent (see Supplementary Table 5 for gene annotation and LD intervals). There was one locus each on chromosomes 20 and 22; two loci each on chromosomes 1, 4, 10 and 19; three loci on chromosome 5; four loci each on chromosomes 12, 13, 14 and 16; six loci each on chromosomes 6, 15, 17 and 18; seven loci each on chromosomes 2 and 3; eight loci on chromosome 7; 12 loci on chromosome 11; and 20 loci on chromosome 8. Five SNPs were exonic, located in *MSRA*, *NOS1*, *PINX1*, *ZCCHC14*, and *C12orf49* genes; a further two were coding SNPs in *RPP21* (a missense mutation) and *AGBL1* (synonymous). For the 116 independent SNPs, evidence of expression quantitative trait loci (eQTL) was explored using the GTEx database and this confirmed that 44 of these were eQTLs (Supplementary Table 5). A Regulome DB score (http://www.regulomedb.org/) was used to identify SNPs with a likely regulatory function. Thirty-three of the 116 SNPs were included in the Regulome DB database and eight of these had a score < 3, indicating that they are likely to bind to DNA and be involved in gene regulation (Supplementary Table 5).

**Figure 1.**
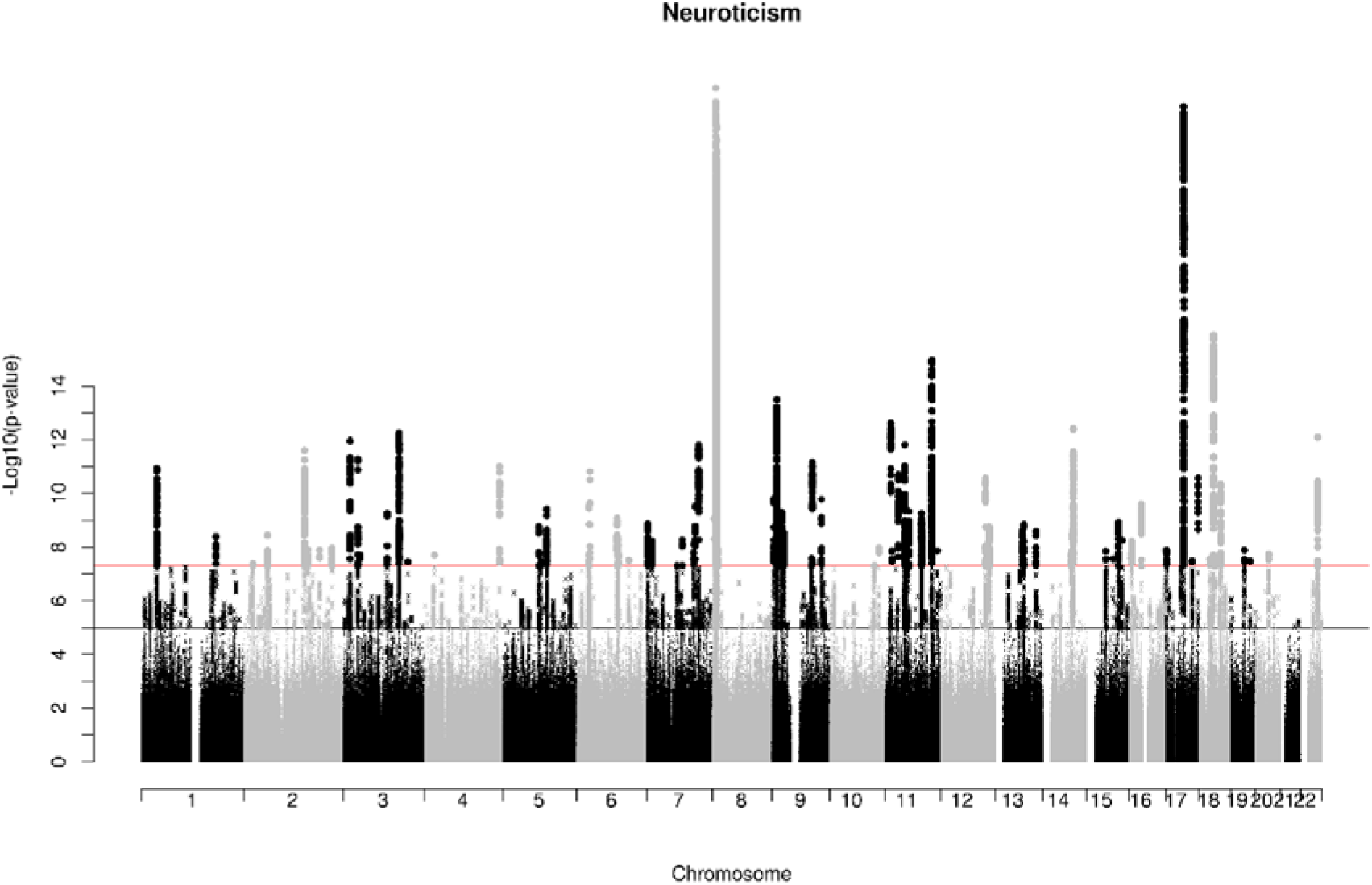
GWA results for neuroticism in 329,821 UK Biobank individuals.

The two SNPs—rs6981523 and rs9611519—previously identified for neuroticism in 23andMe ^16^ were similarly significant in our larger sample, with respective p-values of 4.7 × 10^−22^ and 1.17 × 10^−10^ and consistent direction of allelic effect. SNP rs35855737, significant in the GPC-2 GWA of neuroticism ^17^, was not significant, P = .069. Previous association of SNPs within 8p23.1 linked to an inversion polymorphism (based partly on a subsample of UK Biobank) was even stronger in our study, with the lead SNPs, rs2572431 ^7^ and rs12682352 ^3^, showing respective p-values of 1.33 × 10^−18^ and 1.11 × 10^−24^. Our lead SNP in this region was rs2921036 (P = 8.04 × 10^−26^), an intergenic SNP located within a tighter region of LD (8,092,025-8,863,059 base pairs) within the larger ~4 Mb region (regional association plot is shown in Fig. 2). The 8p23.1 locus was previously cited as important in developmental neuropsychiatric disorders ^18^ which may be relevant for personality traits, which emerge early in life, and could therefore be influenced by the same developmental processes.

**Figure 2.**
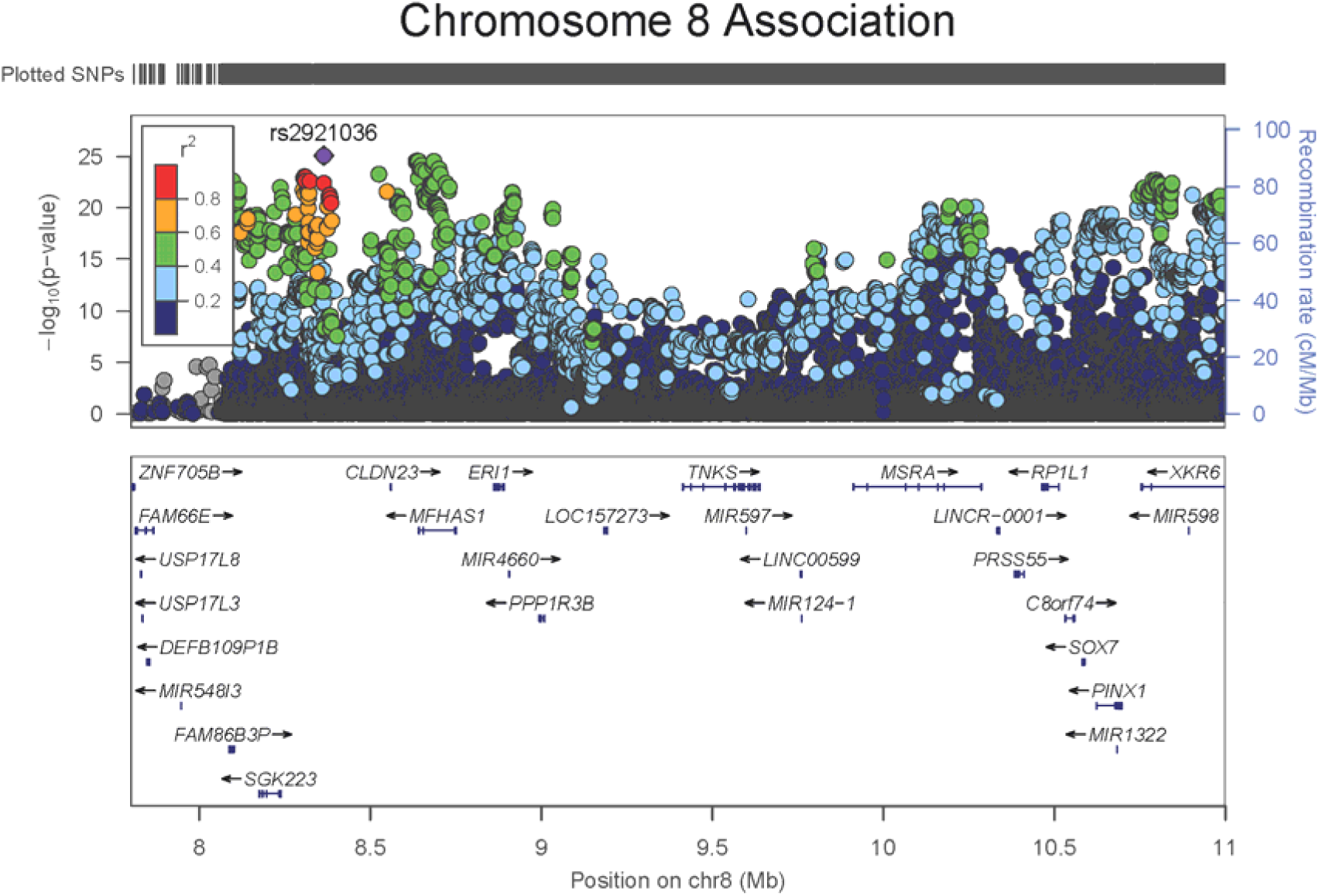
Regional association plot for chromosome 8p suggestive/significant signal.

Replication of the significant association signals in UK Biobank was sought from the results of a GWA meta-analysis of neuroticism which included 23andMe (N = 59,206) ^16^ and the Genetics of Personality Consortium (GPC-2; N = 63,661) ^17^. Of the genome-significant SNPs in UK Biobank, 10,171 were present in the 23andMe and GPC meta-analysis GWA results. Replication was observed for 984 of these at a conservative Bonferroni-corrected level of P < 4.9 × 10^−6^ and all these replicated SNPs increased in significance when meta-analysed with the discovery sample, indicating consistent allelic effect. Supplementary Table 6 shows the effect allele, effect size, and p-value for the discovery, replication, and discovery + replication meta-analysis samples. Of the 116 independent associated SNPs, 111 were present in the replication cohort, with 51 nominally significant (P < .05; see Supplementary Table 5), and 15 at a Bonferroni-corrected level (P < .00045; see Table 1). Figures 3 and 4 show the regional association plot for chromosomes 22 and 11 which, like chromosome 8, showed multiple genes present in the associated LD region. All other replicated independent loci were localised or situated nearby only one or two genes.

**Table 1.**
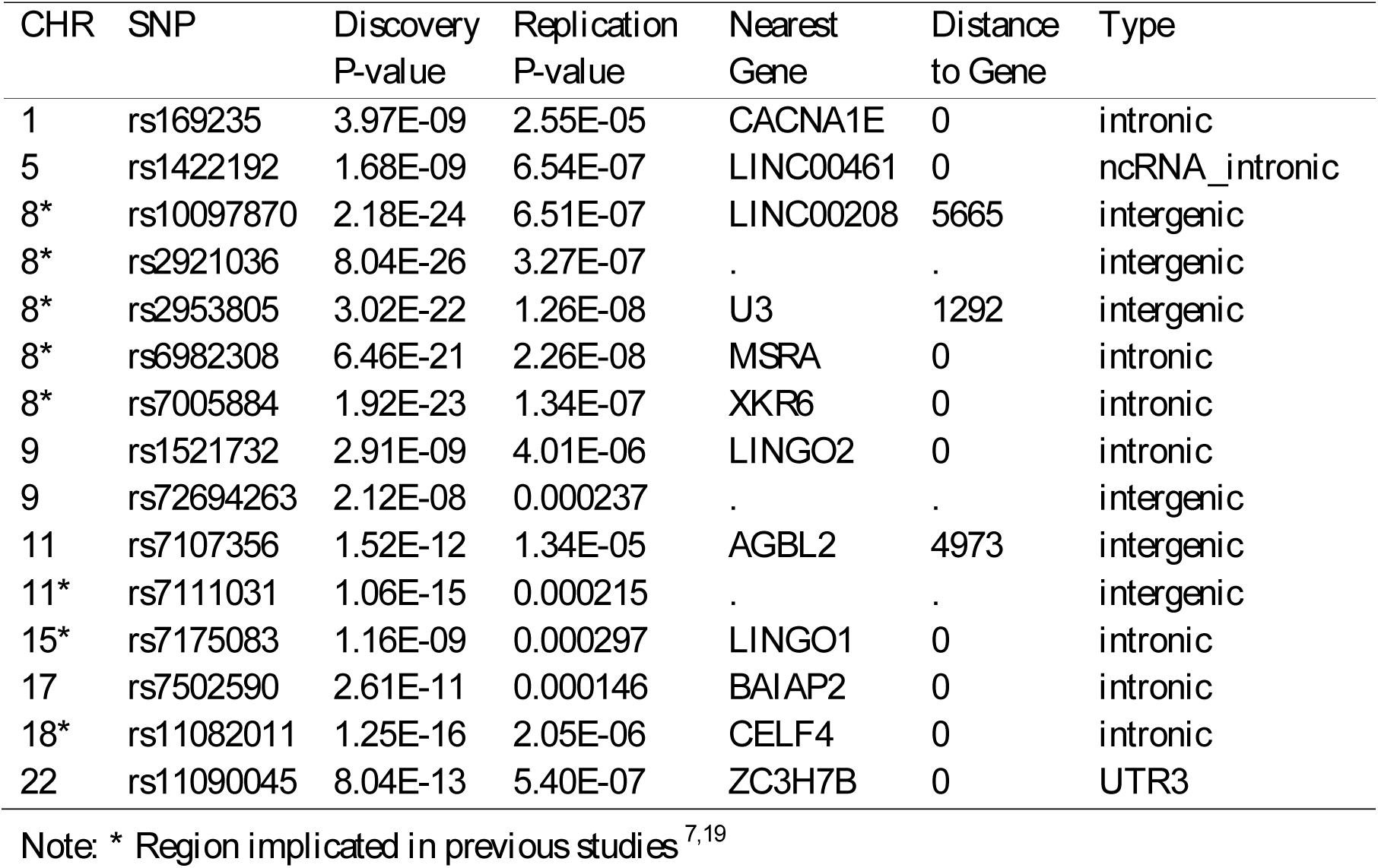
Fifteen independent SNPs associated with neuroticism in UK Biobank most strongly replicated in the meta-analysis of 23andMe and the GPC cohorts.

| CHR | SNP | Discovery<br>P-value | Replication<br>P-value | Nearest<br>Gene | Distance<br>to Gene | Type |
| --- | --- | --- | --- | --- | --- | --- |
| 1 | rs169235 | 3.97E-09 | 2.55E-05 | *CACNA1E* | 0 | intronic |
| 5 | rs1422192 | 1.68E-09 | 6.54E-07 | *LINC00461* | 0 | ncRNA_intronic |
| 8* | rs10097870 | 2.18E-24 | 6.51E-07 | *LINC00208* | 5665 | intergenic |
| 8* | rs2921036 | 8.04E-26 | 3.27E-07 | . | . | intergenic |
| 8* | rs2953805 | 3.02E-22 | 1.26E-08 | *U3* | 1292 | intergenic |
| 8* | rs6982308 | 6.46E-21 | 2.26E-08 | *MSRA* | 0 | intronic |
| 8* | rs7005884 | 1.92E-23 | 1.34E-07 | *XKR6* | 0 | intronic |
| 9 | rs1521732 | 2.91E-09 | 4.01E-06 | *LINGO2* | 0 | intronic |
| 9 | rs72694263 | 2.12E-08 | 0.000237 | . | . | intergenic |
| 11 | rs7107356 | 1.52E-12 | 1.34E-05 | *AGBL2* | 4973 | intergenic |
| 11* | rs7111031 | 1.06E-15 | 0.000215 | . | . | intergenic |
| 15* | rs7175083 | 1.16E-09 | 0.000297 | *LINGO1* | 0 | intronic |
| 17 | rs7502590 | 2.61E-11 | 0.000146 | *BAIAP2* | 0 | intronic |
| 18* | rs11082011 | 1.25E-16 | 2.05E-06 | *CELF4* | 0 | intronic |
| 22 | rs11090045 | 8.04E-13 | 5.40E-07 | *ZC3H7B* | 0 | UTR3 |

Note: \* Region implicated in previous studies <sup>7,19</sup>

**Figure 3.**
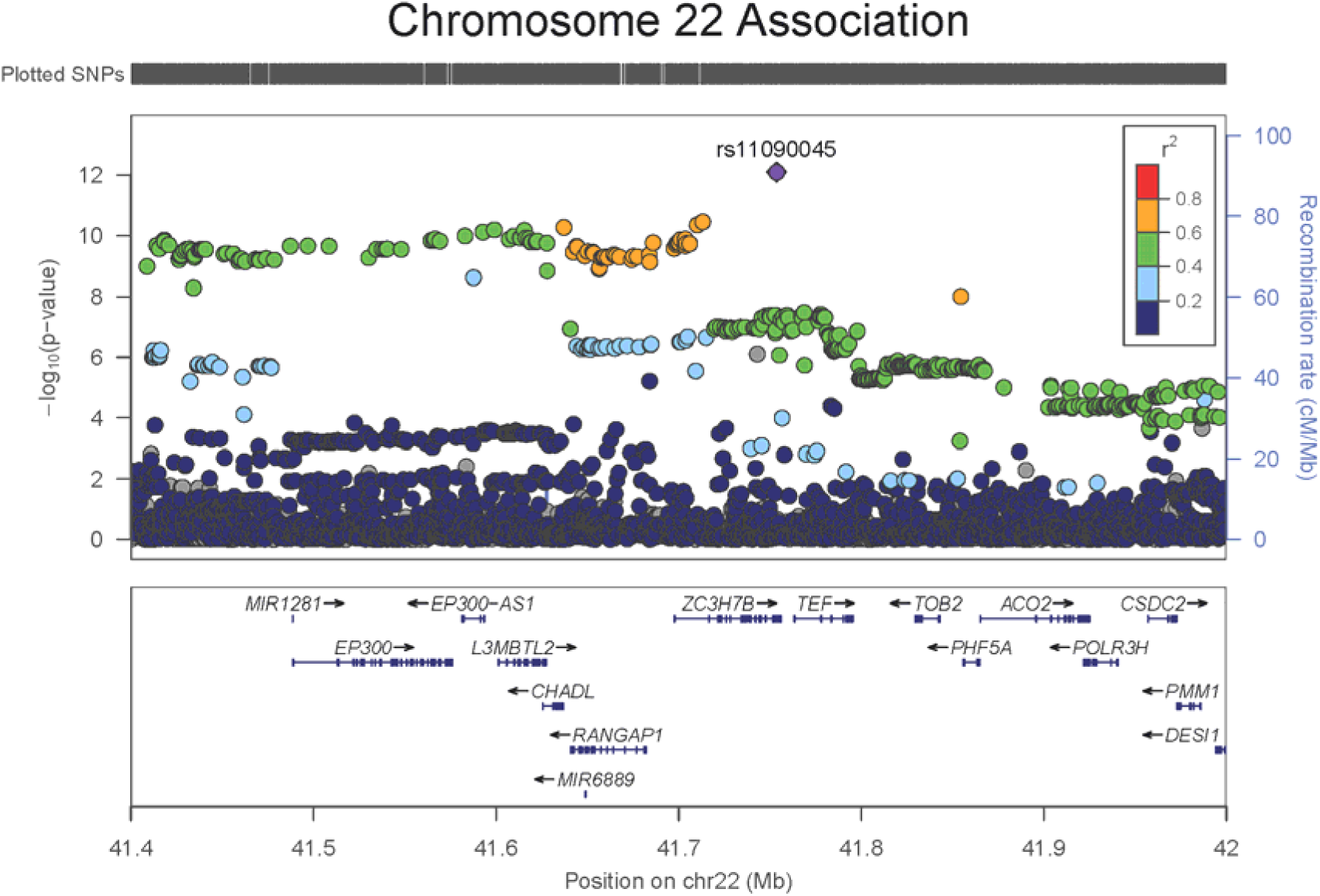
Regional association plot for chromosome 22 suggestive/significant signal.

**Figure 4.**
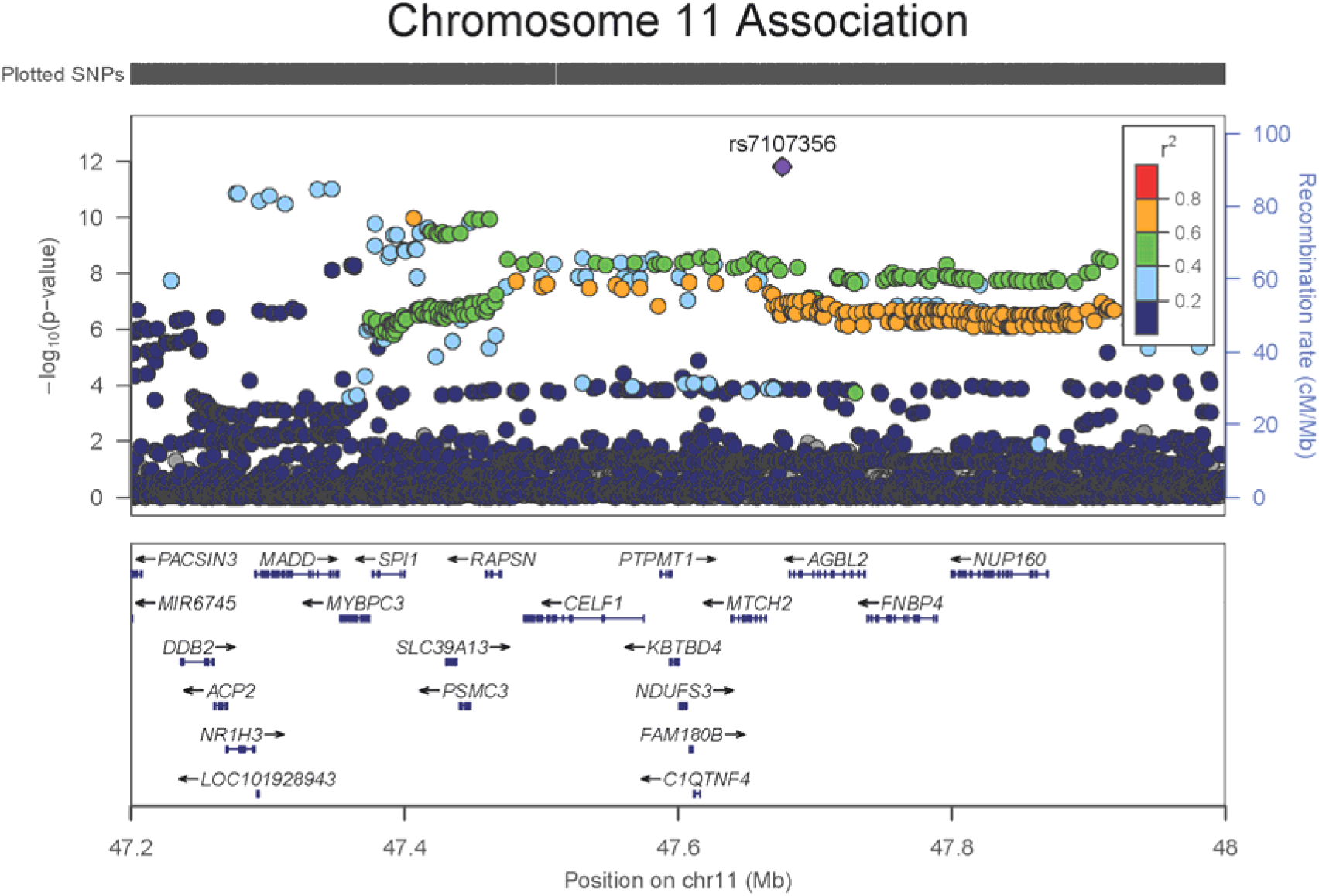
Regional association plot for chromosome 11 suggestive/significant signal.

Gene-based analysis of the GWA results was performed using MAGMA ^20^ ; 249 genes were significantly associated at a Bonferroni-corrected level (α = 0.05 / 18,080; *P* < 2.77 × 10^−6^ ; Supplementary Table 7). These genes corresponded with the genes annotated to the single SNP GWA findings. Three of these were genes (*STH*, *HIST1H3J*, *HIST1H4L*) containing a single SNP. Of the replicated independent GWA SNPs that were in/nearby genes, the following significant genes were corroborated in the gene-based results: *CACNA1E*, *XKR6*, *MSRA*, *LINGO2*, *AGBL2*, *CELF4*, *ZC3H7B* and *BAIAP2*. SNP rs6981523, previously identified in 23andMe, was an intergenic SNP near *XKR6;* this gene was the second most significant gene in our gene-based analysis (P = 6.55 × 10^−32^). *L3MBTL2* and *CHADL*, wherein 23andMe’s other significant SNP, rs9611519, resided, showed respective gene-based p-values of 2.40 × 10^−6^ and 1.15 × 10^−6^.

Pathway analysis in MAGMA highlighted five significant gene ontology pathways (family-wise error P < 1.21 × 10^−6^): neuron spine (cellular), homophilic cell adhesion via plasma membrane adhesion molecules (biological), neuron differentiation (biological), cell cell adhesion via plasma membrane adhesion molecules (biological), and neurogenesis (biological). See Table 2 for further details.

**Table 2.** Significant gene ontology pathways for neuroticism in UK Biobank.

| **Pathway** | **Number of genes** | **Beta** | **SE** | **P-value** | **Corrected P** | **Definition** |
| --- | --- | --- | --- | --- | --- | --- |
| Neuron Spine | 147 | 0.56<br>0 | 0.107 | $7.77 \times 10^{-8}$ | 0.0282 | A small membranous protrusion, often ending in a bulbous head and attached to the neuron by a narrow stalk or neck. |
| Homophilic Cell Adhesion Via Plasma Membrane Adhesion Molecules | 115 | 0.49<br>0 | 0.0938 | $8.81 \times 10^{-8}$ | 0.0289 | The attachment of a plasma membrane adhesion molecule in one cell to an identical molecule in an adjacent cell. |
| Neuron Differentiation | 1341 | 0.14<br>5 | 0.0288 | $2.36 \times 10^{-7}$ | 0.0357 | The process in which a relatively unspecialized cell acquires specialized features of a neuron. |
| Cell Cell Adhesion Via Plasma Membrane Adhesion Molecules | 828 | 0.18<br>3 | 0.0364 | $2.72 \times 10^{-7}$ | 0.0372 | The attachment of one cell to another cell via adhesion molecules that are at least partially embedded in the plasma membrane. |
| Neurogenesis | 195 | 0.41<br>9 | 0.0859 | $5.35 \times 10^{-7}$ | 0.0439 | Generation of cells within the nervous system |

LD score regression ^21^ was used to estimate the genetic correlation between neuroticism and a variety of mental health traits (Supplementary Table 8). The strongest correlation was observed for depressive symptoms (r_g_ = .82, SE = .03). Major depressive disorder and subjective wellbeing showed moderate-to-strong correlations (~.68), schizophrenia, ADHD and anorexia nervosa showed significant and moderate-to-low correlations (~.20), and Alzheimer’s disease had a low correlation (.10). Genetic correlations with ADHD, bipolar disorder and Alzheimer’s disease, which were not significant in a previous study using ~108,000 UK Biobank participants, ^22^ were now significant. The genetic correlation of one between Eysenck neuroticism and other neuroticism scales (used by 23andMe and the GPC) confirms that GWA meta-analysis based on different measurement instruments is valid. Polygenic profile analyses indicated that the neuroticism polygenic score explained 2.75% of the variance in neuroticism (β = .19, P = 1.26 × 10^−4^) and 0.8% of the variance in depression status (OR = 1.24, P = 2.80 × 10^−8^) in Generation Scotland. The results for each threshold can be found in Supplementary Table 9.

Given its strong association with human misery and happiness, its protean predictive power for health—from anxiety and depression to longevity ^5,23^—, and the huge personal and societal burdens it brings, understanding the environmental and genetic origins of people’s differences in neuroticism is a health priority ^6^. The combination, in UK Biobank, of a large sample and a well-validated neuroticism scale has afforded the discovery of 116 genetic loci that influence neuroticism levels, most of them novel. These discoveries promise paths to understand the mechanisms whereby some people become depressed, and of broader human differences in happiness, and they are a resource for those seeking novel drug targets for major depression. As proof-of-principle, the *CRHR1* gene (highlighted in our SNP and gene-based analysis) has been associated with anxiety, depression and neuroticism ^3,24,25^ and is involved in normal hormonal responses to stress; the glucocorticoid pathway is thus a relevant and well-known target. After millennia in which scholars and researchers have sought the sources of individual differences in people’s proneness to dysphoria ^26^, the present study adds significantly to explaining the (genetic) anatomy of melancholy.

## Online Methods

### Genome-wide association analysis in UK Biobank

An imputed dataset, including >92 million variants, referenced to the UK10K haplotype, 1000 Genomes Phase 3, and Haplotype Reference Consortium (HRC) panels was available in UK Biobank. The current analysis includes only those SNPs available in the HRC reference panel. Quality control filters were applied (see online Supplementary Methods) which resulted in 18,485,882 imputed SNPs for analysis in 329,821 individuals. The GWA of neuroticism was conducted using BGENIE ^12^, a program specifically developed to analyse UK Biobank data in a fast and efficient manner. Further information can be found at the following URL: https://jmarchini.org/bgenie/. A linear SNP association model was tested which accounted for genotype uncertainty. Neuroticism was pre-adjusted for age, sex, genotyping batch, genotyping array, assessment centre, and 40 principal components to speed up analysis.

The number of independent signals from the GWAS was determined using LD-clumping in PLINK v1.90b3i ^27^ (**https://www.cog-genomics.org/plink2**). The LD structure was based on SNPs with a p-value < 1 × 10^−3^ that were extracted from the imputed genotypes. Index SNPs were identified (P < 5 × 10^−8^) and clumps were formed for SNPs with P < 1 × 10^−5^ that were in LD (R^2^ > 0.1) and within 500kb of the index SNP. SNPs were assigned to no more than one clump.

### Meta-analysis of GWA Results

Two meta-analyses were performed. To check for replication of the top GWA signals in UK Biobank, results from a meta-analysis of 23andme ^16^ and the Genetics of Personality Consortium (GPC) ^17^ were used. This meta-analysis was conducted using METAL ^28^ and due to the lack of phenotype harmonisation across the cohorts, a sample size weighted metaanalysis was preferred. For all significant SNPs in UK Biobank replicated in the metaanalysis of 23andMe and the GPC, an additional meta-analysis including UK Biobank was performed using the same method.

### Genome-wide Gene-based Analysis

Gene-based analysis of neuroticism was performed using MAGMA ^20^, which provides gene-based statistics derived using the results of the GWA analysis. Genetic variants were assigned to genes based on their position according to the NCBI 37.3 build, with no additional boundary placed around the genes. This resulted in a total of 18,080 genes being analysed. The European panel of the 1000 Genomes data (phase 1, release 3) was used as a reference panel to account for linkage disequilibrium. A genome-wide significance threshold for gene-based associations was calculated using the Bonferroni method (α=0.05/18,080; P < 2.77 × 10^−6^).

### Functional annotation and gene expression

For the 116 independent genome-wide significant SNPs identified by LD clumping, evidence of expression quantitative trait loci (eQTL) and functional annotation were explored using publicly available online resources. The Genotype-Tissue Expression Portal (GTEx) (http://www.gtexportal.org) was used to identify eQTLs associated with the SNPs. Functional annotation was investigated using the Regulome DB database ^29^ (http://www.regulomedb.org/).

### Pathway Analysis

Biological pathway analysis was performed on the gene-based analysis results. This gene-set enrichment analysis was conducted utilising gene-annotation files from the Gene Ontology (GO) Consortium (**http://geneontology.org/)** ^30^ taken from the Molecular Signatures Database (MSigDB) v5.2. The GO consortium includes gene-sets for three ontologies; molecular function, cellular components and biological function. This annotation file consisted of 5,917 gene-sets which were corrected for multiple testing correction using the MAGMA default setting correcting for 10,000 permutations.

### Linkage Disequilibrium Score Regression

Univariate Linkage disequilibrium Score (LDSC) regression ^14^ was used to test for residual stratification in our GWAS summary statistics and to derive a heritability estimate. An LD regression was performed by regressing the GWA test statistics (χ^2^) on to each SNP’s LD score (the sum of squared correlations between the minor allele frequency count of a SNP with the minor allele frequency count of every other SNP). This regression allows for the estimation of heritability from the slope, and a means to detect residual confounders, the intercept. Bivariate LDSC regression ^21^ was used to derive genetic correlations between neuroticism and the following phenotypes: attention deficit hyperactivity disorder (ADHD), Alzheimer’s disease, schizophrenia, anorexia nervosa, depressive symptoms, major depressive disorder, and subjective wellbeing. For Alzheimer’s disease, a 500-kb region surrounding *APOE* was excluded and the analysis re-run (Alzheimer’s disease (500kb)). Further details, including source of GWA summary statistics can be found in the online supplementary material.

### Polygenic Prediction into Generation Scotland

Polygenic profile analyses were performed to predict neuroticism and depression status in Generation Scotland (GS) ^19^. Polygenic profiles were created in PRSice ^31^ using the UK Biobank neuroticism SNP-based association results, for 7,388 unrelated individuals in GS. SNPs with a MAF <0.01 were removed prior to creating the polygenic profiles. Clumping was used to obtain SNPs in linkage disequilibrium with an r^2^ < 0.25 within a 250kb window. Individuals were removed from GS if they had contributed to both UK Biobank and GS (n = 302). Polygenic profile scores were created based on the significance of the association in UK Biobank with the neuroticism phenotype, at p-value thresholds of 0.01, 0.05, 0.1, 0.5 and 1 (all SNPs). Linear regression models were used to examine the associations between the polygenic profile and neuroticism score in GS, adjusting for age at measurement, sex and the first 10 genetic principal components to adjust for population stratification. Logistic regression models were used to examine depression status, adjusting for the same covariates as in the neuroticism models. The false discovery rate (FDR) method was used to correct for multiple testing across the polygenic profiles for neuroticism at all five thresholds ^32^.

### Data Availability

The GWA results generated by this analysis will be made publicly available via the UK Biobank repository.

## Additional information

### URLs

UK Biobank Resource: http://www.ukbiobank.ac.uk

Regulome Database: http://www.regulomedb.org/

PLINK V2: https://www.cog-genomics.org/plink2

Genotype-Tissue Expression Portal: http://www.gtexportal.org

Gene Ontology: http://geneontology.org

## Acknowledgements

This research has been conducted using the UK Biobank Resource (Application Nos. 10279 and 4844). This work was supported by The University of Edinburgh Centre for Cognitive Ageing and Cognitive Epidemiology, part of the cross council Lifelong Health and Wellbeing Initiative (MR/K026992/1); funding from the Biotechnology and Biological Sciences Research Council (BBSRC) and Medical Research Council (MRC) is gratefully acknowledged. This report represents independent research part-funded by the National Institute for Health Research (NIHR) Biomedical Research Centre at South London and Maudsley NHS Foundation Trust and King’s College London. W.D.H. is supported by a grant from Age UK (Disconnected Mind Project). A.M.M. and I.J.D. are supported by funding from a Wellcome Trust Strategic Award (104036/Z/14/Z).

## Author Information

Centre for Cognitive Ageing and Cognitive Epidemiology, Department of Psychology, School of Philosophy, Psychology and Language Sciences, The University of Edinburgh, Edinburgh, UK

Michelle Luciano, Saskia P Hagenaars, Gail Davies, W David Hill, Riccardo E Marioni, Sarah E Harris David C Liewald, Chloe Fawns-Ritchie, Toni-Kim Clarke, Masoud Shirali, Mark J Adams, David M Howard, Andrew M McIntosh, Catharine R. Gale & Ian J Deary

Division of Psychiatry, University of Edinburgh, Edinburgh, UK Toni-Kim Clarke, Masoud Shirali, Mark J Adams, David M Howard & Andrew M McIntosh

Centre for Genomic & Experimental Medicine, MRC Institute of Genetics & Molecular Medicine, University of Edinburgh, Western General Hospital, Edinburgh, UK. Riccardo E Marioni & Sarah E Harris

MRC Social, Genetic and Developmental Psychiatry Centre, Institute of Psychiatry, Psychology & Neuroscience, King’s College London, De Crespigny Park, Denmark Hill, London, UK Saskia P Hagenaars & Cathryn M Lewis

MRC Lifecourse Epidemiology Unit, University of Southampton, Southampton, UK. Catharine R. Gale

## Contributions

M.L. drafted the manuscript with contributions from D.W.H. and I.J.D. G.D., D.C.L., R.E.M., M.J.A. and D.M.H. performed quality control of UK Biobank data and/or Generation Scotland. M.L, G.D, S.P.H., and M.S. analysed the data. T-K.C., C.F-R., and S.E.H. performed/assisted with downstream analysis. M.L. and I.J.D. co-ordinated the work. All authors commented on and approved the manuscript.

## Supplementary

Online Methods and Results

Online Excel Results Tables (large)

